# Subcortical-hippocampal circuits for mediating impaired contextual fear memory after an acute shift of the light/dark phase

**DOI:** 10.64898/2026.07.30.741717

**Authors:** Lara Mariel Chirich Barreira, Hannah Gapp, Anne Albrecht

## Abstract

Acute disturbances of the light-dark cycle may lead to cognitive impairments associated with disturbances in hippocampal functions in humans and rodent models that are potentially governed by subcortical modulation. In this study, we applied a jet-lag-like model in mice by introducing a six-hour delay of the switch towards the light, inactive phase of mice following a contextual fear conditioning training. Phase delay (PD) resulted in a reduced fear memory expression in male but not female mice, associated with an activation of orexinergic neurons in the lateral hypothalamus (LH) as well as of cells in the supramammillary nucleus (SuM) and in the hilus of the dorsal hippocampal dentate gyrus (DG) in male mice, as assessed by immunolabelling for the activity marker c-Fos. Mimicking the overactivation of SuM and DG by chemogenetic stimulation before contextual fear memory retrieval replicated the PD-induced phenotype, suggesting a direct contribution of the SuM and the DG on modulating fear expression after PD. Further circuit analysis by c-Fos revealed a reciprocal interaction between the SuM and the DG. In addition, orexinergic neurons in the LH were activated by chemogenetic stimulation of the SuM. Together, our results reveal that an acute, jet-lag-like phase shift applied during late consolidation stages induced deficits in fear memory expression associated with an overactivation of the SuM-DG pathway and the orexinergic system. These findings may provide insights into the subcortical modulation of memory-relevant circuits, with relevance for acute light-dark rhythm disruptions prevalent in modern societies as well as for disorders associated with memory disturbances.

**Significance statement:** Acute disturbances of the light-dark cycle, as experienced during jet lag or shift work, are increasingly common and can impair memory and cognitive function. Here, we identify a brain circuit underlying jet-lag-induced fear memory deficits that occurs selectively in male, but not female mice. A six-hour delay of the dark phase impaired recall of a previously learned fear memory in males, associated with overactivation of two interconnected brain regions, the supramammillary nucleus and the dentate gyrus of the hippocampus, as well as neurons producing the wake-promoting signal orexin. Artificially mimicking this overactivation was sufficient to reproduce the memory impairment, revealing a hypothalamo-hippocampal circuit that translates circadian disruption into memory deficits in male, with implications for cognitive disorders.

## Introduction

Humans and rodents continuously receive environmental stimuli that must be integrated with their internal state to generate appropriate behavioral responses. Regular rhythmic stimuli, such as the light-dark (LD) cycle, synchronize biological pathways, including those that regulate synaptic plasticity, hormone and neurotransmitter release. This synchronization in turn governs behavioral and physiological processes. In rodent models, the phase of the LD cycle contributes to cognitive performance, such as spatial learning and fear memory (Albrecht & Stork, 2017; Chaudhury & Colwell, 2002; Rawashdeh et al., 2014). In humans, circadian misalignment (e.g., shift work or jet lag) impairs attention, learning, and memory (Özdemir et al., 2013; Vandewalle et al., 2007; Wright et al., 2012). Previous studies have shown that an acute manipulation of the LD cycle after contextual fear condition (CFC) training, either by advancing the light phase (phase advance, PA) or by delaying it (phase delay, PD), affected fear-memory recall 24 h later (Loh et al., 2010). During CFC, individuals learn to associate a neutral multimodal environment with an aversive stimulus. The acute phase shift reduced the fear response to the shock context upon re-exposure (Loh et al., 2010). However, these experiments were performed during the light and not during the dark phase. Moreover, a possible dysregulation of memory-relevant circuits by such acute phase shifts is rather unexplored.

While the hippocampal formation is a key structure for the formation and retrieval of memories, recent evidence suggests a modulation by subcortical inputs via the supramammillary nucleus (SuM) (Vertes, 2015). The SuM projects unidirectionally to the dentate gyrus (DG), to CA2, and to interneurons in the CA1/CA3 regions (Chen et al., 2020; Hayakawa & Zyo, 1996; Y. Li et al., 2020; Tabuchi et al., 2022). SuM terminals co-release glutamate and GABA, thereby modulating hippocampal circuits (Boulland et al., 2008; Hashimotodani et al., 2018; Tabuchi et al., 2022). Interestingly, a medial (SuMM) and lateral (SuML) sub-region can be defined, which preferentially project to the ventral versus dorsal DG (dDG) and CA1/CA3 regions, respectively, differing in their neurotransmitter content (Chen et al., 2020; Hashimotodani et al., 2018; Hayakawa & Zyo, 1996; Y. Li et al., 2020; Vertes, 2015).

The SuM is reciprocally connected with the lateral hypothalamus (LH). SuM neurons projecting to the dDG receive inputs from the LH (Luo et al., 2025), thereby defining a putative LH-SuM-DG pathway. One of the major neuropeptides expressed in the LH is orexin (hypocretin), either as orexin-A (Ox-A) or orexin-B, which derive from the precursor pre-pro-orexin, encoded by an unique gene (De Lecea et al., 1998). The activity of orexinergic neurons in the LH is increased during the active phase and decreased during sleep (Azeez et al., 2018; España et al., 2003; Furlong et al., 2009). Orexinergic neurons project widely throughout the brain and bind to orexin receptors type 1 and 2 (OxR1 and OxR2), thereby taking part in emotion regulation, wakefulness, arousal, anxiety, and memory in rodents (Chung et al., 2014; España et al., 2003; J. Li et al., 2014; Peyron et al., 1998; Shaw et al., 2017). Both receptors are expressed in the hippocampus and the SuM (Krause et al., 2024; Tsuneoka & Funato, 2024), making orexinergic neurons an interesting modulator of the SuM-DG pathway.

In this study, we hypothesize that an acute “jet-lag”-like phase shift of the mice’s active phase impairs fear memory mediated by a LH_orexin_-SuM-DG pathway. Indeed, we found that the expression of contextual fear memory was impaired in male, but not female mice that experienced an acute light shift with a delay of their active phase after footshock exposure. These behavioral changes were associated with an increased activation of orexinergic neurons in the LH as well as of cells in the SuM and the dDG hilus. Mimicking PD-induced overactivation of the DG or SuM by chemogenetic manipulation replicated the behavioral phenotype and supported the involvement of SuM-LH interactions.

## Material and Methods

### Animals

Young adult male and female C57BL/6J mice (9-14 weeks of age) were bred and raised at the central animal facility of the Otto-von-Guericke University Magdeburg. They were kept in groups of two to five individuals per cage, with water and food ad libitum. Standard rearing conditions consisted of a 12/12 h inverse LD cycle (lights on at 8 pm, lights off at 8 am); the phase shift experiments differed from this schedule (see below). All behavioral experiments were performed during the dark, active phase of the animals. All housing and experimental procedures complied with European and German regulations and were approved by the responsible authorities in accordance with German law, namely the Landesverwaltungsamt Saxony-Anhalt (Permission Nr. 42502-2-1717) and its ethical committee. Reporting followed the ARRIVE guidelines.

### Experimental design

#### Contextual fear conditioning (CFC)

CFC was conducted in a fear-conditioning box (26 x 26 x 35 cm, width x depth x height) made of acrylic glass, placed inside a sound-proofed cubicle (Ugo Basile SRL, Gemonio VA, Italy), using a four-day protocol (Fig. 1A). Animals were habituated to the conditioning box in two sessions on two consecutive days by placing the mice in a neutral context (three walls covered with plastic sheets in black, black-white-striped, and black-white-chequered patterns; rough-surfaced floor; cleaned with 0.2% acetic acid). On day 3, training took place in the shock context (transparent walls, grid floor, cleaned with 70% ethanol). Mice were first placed in the shock context for 2 min (pre-training phase), followed by the application of three footshocks (0.4 mA, 1 s each, 20 s intervals). Mice remained in the shock context for an additional 2 min following the last footshock (post-shock phase). The retrieval test was performed the next day by re-exposing the animals to the shock context for 6 min. Throughout all phases, movement was recorded with a camera positioned above the fear-conditioning box (Basler ace acA1300-60gm, Basler, Ahrensburg, Germany) under infrared illumination and analysed automatically using EthoVision video-tracking software (versions 16 and 17; Noldus Information Technology, Wageningen, Netherlands). Freezing, as an expression of defensive behaviour, during the first 2 min of retrieval was quantified as the percentage of time a mouse spent with a mobility below 4% (Pham et al., 2008), corresponding to no more than a 4% change in the pixels of the detected mouse between frames.

**Figure 1:**
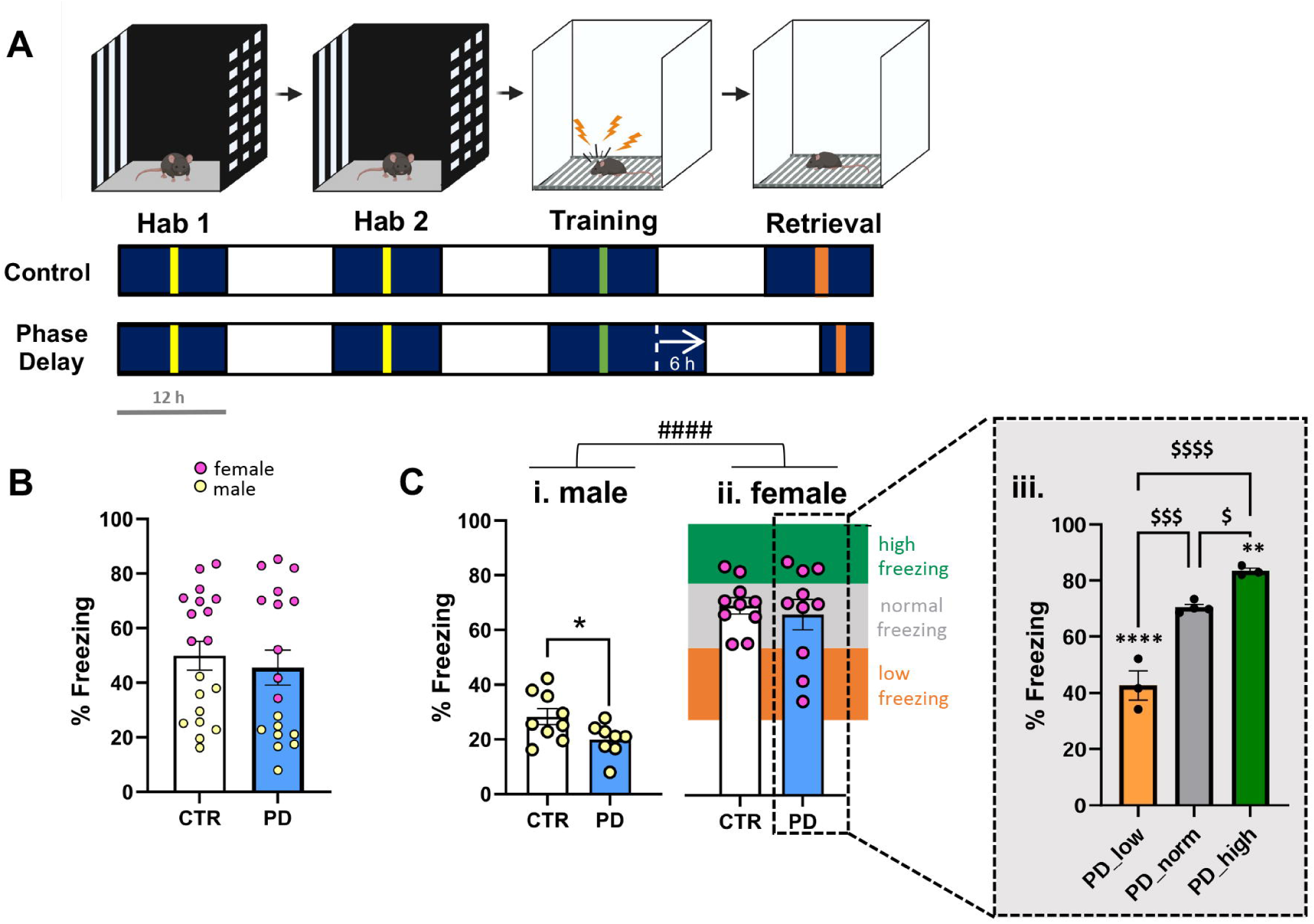
Acute phase delay (PD) after contextual fear conditioning (CFC) training reduces freezing in male mice. **(A)** Schematic of CFC protocol and PD. Habituation 1 and 2 (Hab 1, Hab 2; yellow lines) were conducted in a modified fear-conditioning (FC) box (neutral context) during the dark, active phase of the mice (dark blue bars). Training (green line) took place 24 h later in the original FC box (shock context), where three footshocks were delivered. Control animals remained on a standard 12/12 h light-dark cycle, whereas the PD group experienced a 6 h PD of the dark active phase at the end of training day, followed by a 12 h light phase (white bars). Fear memory was tested during the retrieval session (orange line) in the subsequent dark phase by re-exposure to the shock context. **(B)** PD treatment did not alter overall freezing levels during retrieval when considering animals of both sexes in each group. (C) Main effects of sex and PD were discovered on freezing levels, with females showing higher freezing. Post hoc test and separate comparisons by sex showed that PD reduced freezing in male (i), but not female mice (ii). Considering the higher variance of freezing reponses after PD in female mice, a behavioral profiling approach was used to define subgroups relative to the mean freezing response of the control groups (low-freezing: PD_low, orange bar, below the 20^th^ percentile; normal-freezing: PD_norm, grey bar, > 20^th^ & <80^th^ percentile; high-freezing: PD_high, green bar, above the 80^th^ percentile). Freezing levels of the PD_high and PD_low groups differed significantly from the female CTR group and from the PD_norm subgroup (iii). Data presented as individual data points and mean ± SEM. * significant difference CTR vs. PD, p<0.05; #### main effect of sex p<0.0001; & significant differences PD subgroups vs. CTR, p<0.01; &&& p<0.001; $ significant differences between PD subgroups, p<0.05; $$$ p<0.001; $$$$ p<0.0001.

#### Acute phase shifts of the light-dark cycle after contextual fear conditioning

To produce an acute light-shift, male and female mice were transferred to an airflow cabinet (Uni-Protect 1603, EHRET Labor und Pharmatechnik GmbH & Co. KG, Emmendingen, Germany), which had been modified with two individually illuminated compartments. The temperature was maintained at 21.5 °C and relative humidity at 40-60%, matching the main housing room conditions. Mice were transferred to the cabinet one week before the start of the behavioral experiments to adapt to the new environment and were kept on an inverse 12/12 h LD cycle. The onset of the light phase was defined as Zeitgeber Time 0 (ZT0). For all animals, habituation sessions and CFC training took place around ZT 17-18 (5-6 h after the onset of the dark phase).

After the training session, the control group continued on the standard 12/12 h LD cycle, whereas the experimental group was subjected to a six-hour phase shift: either a phase delay (PD), where the active, dark phase of the animals was prolonged (see Fig. 1A), or a phase advance (PA) in male mice, in which the active, dark phase ended 6 h earlier, around one hour after the end of CFC training (see Fig. S1). Following the shift, a 12-h light period was maintained for all groups. The retrieval of the fear memory was performed the following day during the dark phase, between ZT 17-18 in the control and PA groups, and between new ZT 13-14 for the PD group (1 h after lights-off). 60 min after memory retrieval, transcardial perfusion was performed with 4% paraformaldehyde (PFA) for tissue fixation and subsequent immunohistochemical analysis.

For ***chemogenetic DG and SUM activation experiments***, a standard 12/12 h LD cycle was maintained throughout the CFC experiments, with standard housing of all animals in individually ventilated cages. All manipulations were performed during the dark phase (habituation: ZT 13-18; training: ZT 14-17; retrieval: ZT 15-18).

### Injection of viral vectors

Mice were placed in a stereotactic frame (World Precision Instruments, Friedberg, Germany) under isoflurane inhalation anesthesia (1.5-2.5 Vol% in O_2_) and analgesic control with carprofen (5 mg/kg, i.p.). A beveled 33G needle mounted on a 10µl NanoFil microliter syringe (World Precision Instruments, Friedberg, Germany) was placed in the drill holes and lowered to either reach the dDG or the SuM at the following coordinates relative to Bregma (Nishijima et al., 2007): DG: -2.0 mm AP, ± 1.3 mm ML and -2.0 DV; SuM: -2,8 mm AP, ± 0.5 mm ML, and -5,2 mm DV. Animals were bilaterally injected with either a control vector, pAAV-hsyn-EGFP (a gift from Brian Roth; Addgene viral prep #50465-AAV8; http://n2t.net/addgene:50465; RRID:Addgene_50465) titer 1.6x10^13^, or an activating chemogenetic vector pAAV-hsyn-hM3D(Gq)-mcherry (a gift from Brian Roth; Addgene viral prep #50474-AAV8; http://n2t.net/addgene:50474; RRID:Addgene_50474) titer 2.4x10^13^, using a MICRO-2T microinjection syringe pump (World Precision Instruments) at a rate of 100 nl/min and with an injection volume of 1 µl/ hemisphere for DG and 200 nl/hemisphere for SuM. The needle remained in situ for 5 min prior to withdrawal. Fear conditioning experiments commenced after at least a 14-day recovery phase in the home cage.

***Chemogenetic activation of the DG or the SuM*** was induced by injecting CNO (DG: 1 mg/kg i.p.; SuM: 5 mg/kg i.p.) 45 min prior to the retrieval test (Fig. 5A and 7A). For the DG-injected cohort, perfusion was carried out 30 min after the retrieval phase for subsequent RNAscope analysis, and for the SuM-injected cohort, 60 min after retrieval for immunohistochemistry (IHC) analysis and verification of viral expression sites. ***Viral expression*** was checked in 3-4 alternating sections of the dorsal hippocampus or the SuM by imaging GFP and mCherry fluorescence with a Leica Thunder Dmi8 microscope (Leica, Wetzlar, Germany). Only mice with correct bilateral injection sites were included into the study.

### Immunohistochemistry

For analysis of neuronal activation, c-Fos IHC was performed as described before (Albrecht et al., 2022). After transcardial perfusion, brains were removed immediately and post-fixed in 4% PFA at 4°C overnight, followed by an immersion in 30% sucrose for cryoprotection, snap freezing of the tissue in 2-methylbutane at −55°C and storage at −80°C.

The brains were sectioned using a cryostat microtome (Leica CM1900) in series of 30 µm thick coronal slices at the level of the dorsal hippocampus (AP: -1.58 to -2.06 mm), the SuM (AP: -2.70 to -3.08 mm), and the LH (AP: -1.34 to -1.94 mm from Bregma).

Free-floating sections were washed in 0.1 M PB and incubated for 1 h at room temperature (RT) in a blocking solution containing 5% normal goat serum (NGS) and 5% bovine serum albumin (BSA) in 0.1 M Phosphate buffer + 0.3% Triton (PBT).

For single labeling of c-Fos, sections were incubated with rabbit anti-c-Fos antibody (1:500; Cell Signaling #2250, Danvers, MA, USA) for 48 h at 4°C, followed by application of biotinylated goat anti-rabbit secondary antibody (1:200; Jackson ImmunoResearch, Ely, UK) in PBT and later labelling with Cy2, Cy3 or Cy5 via Streptavidin (1:1000; Jackson ImmunoResearch, Ely, UK) in 0.1 M PB.

For double labeling of c-Fos and Ox-A in the LH, the protocol was adapted by using a mixture of rabbit anti-c-Fos (1:1000; Cell Signaling #2250) and mouse anti-orexin-A primary antibodies (1:500; Santa Cruz #sc-80263). As secondary antibodies, biotinylated goat anti-rabbit (1:200; Jackson ImmunoResearch, Ely, UK) and goat anti-mouse Alexa Fluor 594 (1:1000; Abcam ab150080) were added for 2 h at RT. Nuclei were visualized with DAPI (300 nM; Thermofischer, Waltham, MA, USA).

Images were acquired using a Leica Thunder DMi8 epifluorescence microscope (Leica, Wetzlar, Germany). Both hemispheres were imaged from at least two sections per animal and region. Animals were excluded from analysis if fewer than two sections sufficiently represented the target brain area. Using ImageJ software (version 1.53t; Java 1.8.0_322), regions of interest (ROIs) were outlined for each brain area and cell counting was performed manually within the respective ROIs by an experimenter blinded to the experimental groups. The cell number was normalized to ROI area and averaged across sections and left and right hemispheres for each individual animal. The resulting mean cell density values and the percentage of c-Fos/Ox-A double-positive cells per animal were used for further statistical comparison.

### Fluorescence in situ hybridization (RNAscope)

To allow for further analysis of SuM activity and specifically of OxR1 mRNA-positive SuM neurons, fluorescence in situ hybridization was performed using the RNAscope Multiplex Fluorescent Assay Version 2 (#323100) for fixed frozen brain tissue (Advanced Cell Diagnostics, Newark, USA) in a subset of animals (selected for freezing levels representative for each group). The assay was performed according to the manufacturer’s instructions and as described previously (Krause et al., 2024) in brains perfused 30 min after contextual fear memory retrieval following chemogenetic activation of the DG. In brief, after rinsing slices with 1 M phosphate-buffered saline (PBS), pre-treatment by baking for 30 min at 60°C (dry environment), post-fixation in 4% PFA for 15 min at 4°C, and dehydration by successive steps at increasing ethanol concentrations at RT, slices were incubated for 15 min at 98°C in target retrieval solution, washed, and rinsed with 100% ethanol. Slices were then hybridized for 2 h with the probes of interest at a dilution of 1:50 for Channel 3 probes into the Channel 1 probe solution: c-Fos as channel 1 probe (Mm-Fos, catalog #316921) in the dorsal hippocampus; c-Fos as channel 1 probe (Mm-Fos, catalog #316921) and orexin as channel 3 probe (Mm-Hcrt, catalog #490461) for the LH; OxR1 as channel 1 probe (Mm-Hcrtr1, catalog #466631) and c-Fos as channel 3 probe (Mm-Fos, catalog #316921) in the SuM. Additional slices were incubated with negative or positive control probes (RNAscope 3-plex: Positive Control Probe #320881; Negative Control Probe, #320871; ACD, Newark, USA). Following the hybridization step, samples were washed with fresh buffer and incubated overnight in 5× SSC at RT, until hybridization with amplifiers 1-3 commenced the next day sequentially. All sections were then incubated first with HRP-C1 (channel 1) or HRP-C3 (channel 3) and then with corresponding fluorophores: HRP-C1 and TSA-520 at a 1:1500 for dorsal hippocampus; HRP-C1 with TSA-570 at 1:1500 and HRP-C3 with TSA-520 at 1:24000 for LH; HRP-C1 with TSA-650 at a 1:1500 and HRP-C3 with TSA-520 at 1:1500 for SuM. After a final blocking step of 15 min, slices were incubated with DAPI for 30 s.

Slices were imaged with the Leica Thunder epifluorescence microscope using 2-3 slices per animal. Due to the high c-Fos activation in the dorsal hippocampus following chemogenetic activation of the DG, ***intensity-based quantification*** was performed in this region by normalizing the c-Fos mRNA signal intensity to a background signal, measured in a circular region (∼500-1000 µm²) adjacent to the DG (c-Fos intensity = mean intensity ROI/ mean intensity background). For each animal, intensities were averaged across slices and hemispheres. ***For LH and SuM, a cell-based quantification*** was performed using the Cell Detection Tool (QuPath v0.2.3-v0.3.2; (Bankhead et al., 2017) with the following settings: requested pixel size, 0.5 μm; background radius, 8 μm; median filter radius, 0 μm; sigma, 1.5 μm; minimum area, 20 μm²; maximum area, 400 μm²; and soma extension, 5 μm). Cell detection was manually corrected when necessary, and the intensity threshold for cell recognition was adjusted between 10 and 30. When at least one defined dot was located within the cell soma boundaries determined in this way, the cell was considered positive.

### Statistical analysis approach

Statistical analysis and graph generation were performed using GraphPad Prism for Windows (version 9; GraphPad Software, San Diego, CA, USA). Outliers were identified and excluded by the ROUT method (robust regression and outlier removal) provided by GraphPad Prism Software (Q=1%). This led to the exclusion of one data point for the male control group for c-Fos-positive cells in the pyramidal layer of CA1 in the PD experiment.

In our initial exploratory study to investigate the impact of PD, we first compared control and PD-exposed groups using either an unpaired t-test or, in cases of non-normally distributed data, a Mann-Whitney test. In cases of unequal variances, Welch’s correction was applied. Normality assumptions were tested using the Shapiro-Wilk test, and equality of variances was tested with Levene’s test. Taking into account potential sex differences, two-way ANOVAs were performed for the main effects of sex, PD and interactions. Here, non-normally distributed continuous variables were transformed using a log10 function prior to analysis to satisfy normality assumptions. Sample sizes were determined before with power analysis by the G*Power software using large effect sizes. For a two-way ANOVA for group and sex effects as well as their interactions, an effect size of *f*=0.45 with *a* =0.05 would require ∼ 10 animals per group. Thus, the group sizes used in our exploratory analysis allow to detect large sex effects. Indeed, on the behavioural level, the two-way ANOVA found main effects of PD and sex, indicative of differential responses to the task and its modulation by PD in males and females. Therefore, PD effects were analysed in male and female subgroups separately, which is of particular relevance for the subsequent analysis of relevant circuit activation by cFos IHC. Prior power analysis confirmed that a power of > 80% can be achieved with 8-10 animals per group, as stated before (Botterill et al., 2021). We acknowledge, that in the female control group less than 8 animals were analysed for activation of orexinergic neurons in the LH and of cells in the SUM and dorsal hippocampus. However, since a PD effect was lacking in female mice, we followed a different approach to link freezing behavior and circuit activation in females: Since female mice showed a heterogenous response, we appled a behavioral profiling approach that considers extreme behaviors compared to the control group (Ardi et al., 2016): The mean freezing levels of the female control group were used to define a low freezer subgroup (% freezing < 20^th^ percentile, “PD_low”), a high freezer subgroup (% freezing > 80th percentiles of control group freezing; “PD_high”) and a normal freezing subgroup (% freezing >20/< 80th percentiles of control group; “PD_norm”). The number of PD-exposed female mice falling into these subgroups was low (n=3-4), which impacts statistical power. However, our analysis shows that the profiling approach builds behaviorally relevant subgroups that demonstrate significant effects in the one-way ANOVA that was performed together with a Tukey’s post hoc to compare multiple groups, suggesting sufficient power to analyse large circuit differences between these freezing subgroups.

For the statistical evaluation of the confirmatory experiments in male mice using chemogentic activation of the SuM and DG, either an unpaired t-test with Welch’s correction if required or a Mann-Whitney test for non-normally distributed data was performed. In cases of unequal variances, was applied. Differences were considered significant with a p-value of p<0.05.

## Results

### Acute phase delay reduces fear memory expression in male, but not female mice

To induce a “jet-lag” in mice, the dark phase was acutely prolonged for six hours after CFC training in male and female mice, and fear memory was tested the next day (Fig. 1A). A Mann-Whitney test across the full cohort of animals revealed no significant differences in fear expression via freezing levels (Fig. 1B; n=19 CTR, n=18 PD; Mann-Whitney U-test: U=137, p=0.2779), while a two-way ANOVA revealed significant main effects of sex and PD (Fig. 1Ci+ii; Males: n=9 CTR, n=8 PD; Females: n=10 CTR, n=10 PD; Two-way-ANOVA on log10-transformed data: sex: F(1,33)=121.3, p<0.0001; PD: F(1,33)=5.179, p=0.0295; Interaction: F(1,33)=2.169, p=0.1503). Post hoc Tukeýs test showed that the PD effect was driven primarily by males, who showed a reduction in freezing by PD during re-exposure to the shock context in the following dark phase (Male CTR vs. PD: p=0.0157). In female mice, no PD-induced changes in freezing duration were observed (Female CTR vs. PD: p=0.5572), although their overall freezing levels were higher. Moreover, female mice showed a diverse response to PD, as revealed by behavioral profiling (Ardi et al., 2016). Based on their freezing levels relative to the mean freezing response of the control group, we could identify low-freezing (PD_low; n=3), normal-freezing (PD_norm; n=4), and high-freezing (PD_high; n=3) individuals. Accordingly, PD_high and PD_low groups differed significantly from the control group and from the PD_norm group during contextual fear memory retrieval (Fig. 1Ciii; one-way ANOVA: F(3,12)=45.23, p<0001; Tukey’s post hoc test).

Phase advance paradigm (PA), where the light phase started 6 h earlier after CFC training, did not induce significant changes in the expression of fear in male mice when the recall was tested during the next dark phase after training (Fig. S1B; Student’s T-test: T(6)=0.302, p=0.773).

Together, a sex-specific impact of an acute prolongation of the dark phase on fear memory expression was observed.

### Acute phase delay activates orexinergic neurons in the lateral hypothalamus and the supramammillary-dentate gyrus circuit in male mice

To analyze the activation of the SuM-DG circuit and a possible modulation by orexinergic neurons as regulators of the sleep-wake cycle, c-Fos IHC was performed in brain slices from male and female mice after fear memory retrieval, with and without PD. Given the sex dimorphism observed in behavioral freezing, we evaluated region-specific PD effects using planned within-sex comparisons alongside two-way ANOVA for sex and PD interactions and addressed neuronal activation in females in the different freezing subgroups (see also Fig. 1Ciii).

In the LH, PD did not modify c-Fos-positive density when sexes were pooled (Fig. 2Bi; n=14 CTR, n=17 PD, [one male CTR and one female PD excluded due to insufficient representation of LH]; Unpaired t-test with Welch’s correction T-test: T(28.86)=1.649, p=0.1100). Two-way ANOVA revealed a tendency of effect of sex (Fig. 2Bii+iii; Male: n=8 CTR, n=8 PD; Female: n=6 CTR, n=9 PD; two-way ANOVA: sex: F(1,27)=4.088, p=0.0532; PD: F(1,27)=3.535, p=0.0709; Interaction: F(1,27)=1.325, p=0.2597). The separate analysis per sex confirmed an increased number of c-Fos-positive cells in males after PD (Fig. 2Bii; n=8 CTR, n=8 PD; Mann-Whitney U-test: U=10, p=0.021), but not female mice (Fig. 2Biii; n=6 CTR, n=9 PD; Mann-Whitney U-test: U=25, p=0.864). No differences were observed when considering LH activation across female freezing subgroups (Fig. 2Biv; one-way ANOVA: F(2,6)=0.804, p=0.491). Similarly, while no general PD effect was observed on OxA-positive cell density (Fig. 2Ci; n=14 CTR, n=17 PD; Student’s T-test: T(29)=1.246; p=0.2226), the two-way ANOVA revealed main effects of sex and PD (Fig. 2Cii+iii; Male: n=8 CTR, n=8 PD; Female: n=6 CTR, n=9 PD; two-way ANOVA: sex: F(1,33)=121.3, p<0.0001; PD: F(1,28)=2.406 p=0.1321; interaction: F(1,27)=0.7718, p=0.3874). The sex-specific comparisons showed that the number of OxA positive neurons was increased in PD-exposed male (Fig. 2Cii; n=8 CTR, n=8 PD; Student’s T-test: T(14)=3.785, p=0.002), but not female mice (Fig. 2Ciii; n=6 CTR, n=9 PD; Student’s T-test: T(13)= 0.965, p=0.352). No differences were observed between female freezing subgroups (Fig. 2Civ; one-way ANOVA: F(2,6)=1.349, p=0.328). To determine the activation of specifically orexinergic cells in the LH, c-Fos and OxA co-expression was evaluated. PD-dependent effects were not detected between groups with sexes pooled (Fig. 2Di; n=14 CTR, n=17 PD; Student’s T-test: T(29)=1.246; p=0.2226), but the two-way ANOVA revealed main effects of sex and PD (Fig. 2Dii+iii; Male: n=8 CTR, n=8 PD; Female: n=6 CTR, n=9 PD; two-way ANOVA: sex: F(1,27)=0.52.18, p<0.0001; PD: F(1,27)=7.687, p=0.0100; interaction: F(1,27)=0.6258, p=0.4358). The sex-specific comparisons demonstrated a significant increase in activated OxA neurons in male (Fig. 2Dii; n=8 CTR, n=8 PD; Student’s T-test: T(14)=3.808, p=0.002), but not female mice (Fig. 2Diii; n=6 CTR, n=9 PD; Mann-Whitney U-test: U=20, p=0.456) or between female freezing subgroups (Fig. 2Dvi; one-way ANOVA: F(2,6)=0.580, p=0.588). This suggests an overall increased activation of orexinergic neurons in the LH by PD in male mice.

**Figure 2:**
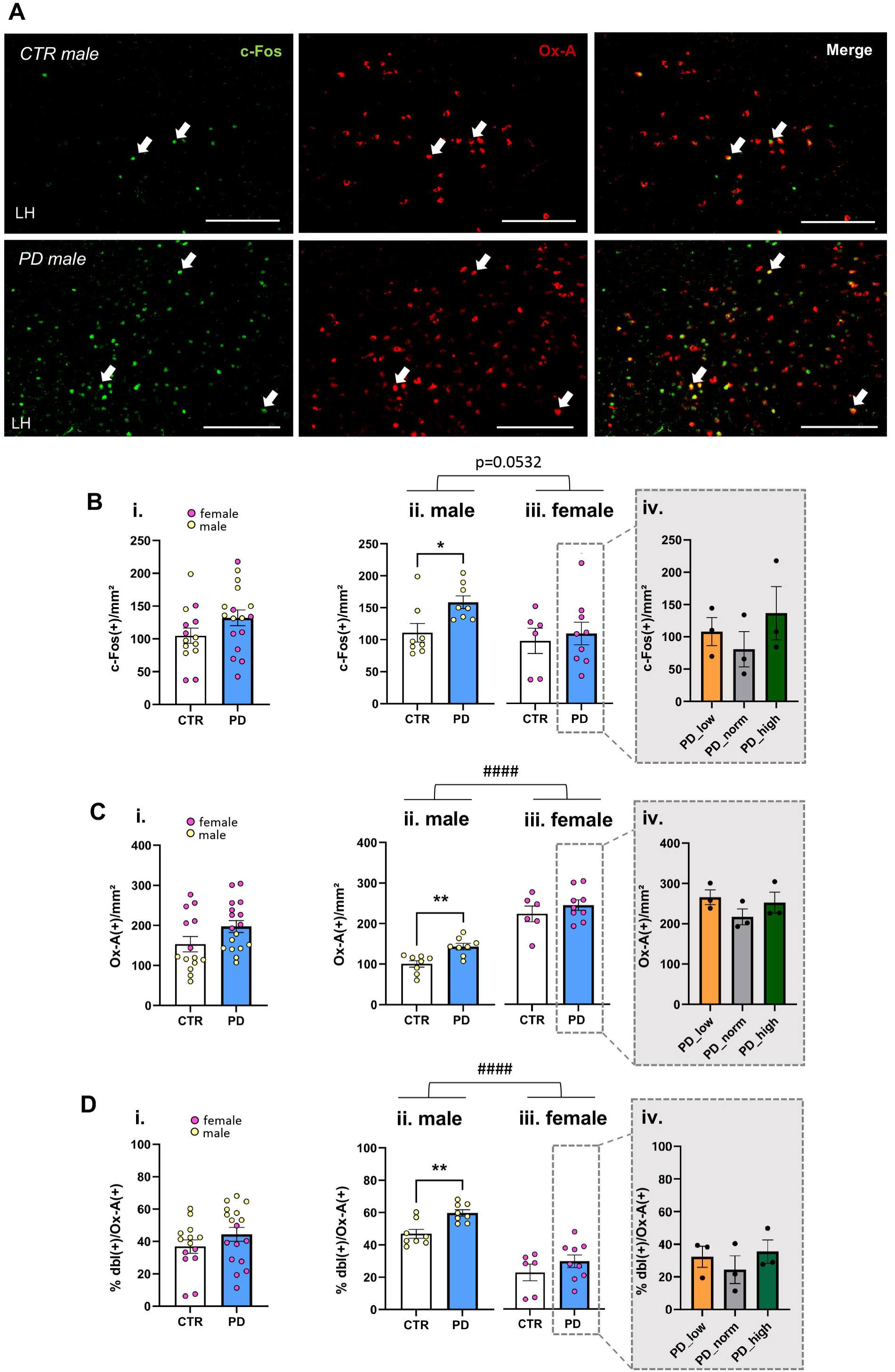
Acute phase delay (PD) activates orexinergic neurons in the lateral hypothalamus (LH) of male mice. **(A)** Representative microphotographs of c-Fos (green, arrow), orexin-A (Ox-A, red, arrow), and double immunolabelling (yellow, arrow in merged panel) in the LH of male mice under control (CTR) and phase delay (PD) conditions, scale bar: 200 µm. **(B)** PD treatment did not alter overall c-Fos(+) cell density in the LH, pooling animals of both sexes (i). Separate analysis per sex revealed that PD increased the number of c-Fos(+) cells in male (ii) but not in female mice (iii). No differences were observed between female PD freezing subgroups (vi). **(C)** PD did not alter the number of OxA(+) cells in the LH when pooling animals of both sexes (i), but two-way ANOVA revealed a main effect of sex. The separate analysis per sex revealed that PD increased the number of Ox-A(+) cells in male (ii), but not female mice (iii). No differences were observed between female PD freezing subgroups (vi). **(D)** Activated orexinergic cells did not differ after PD when considering animals of both sexes (i), but two-way ANOVA revealed a main effect of sex. The separate analysis per sex demonstrated an increased c-Fos-activation of Ox-A(+) cells in male (ii), but not female mice after PD (iii). No differences were observed between female PD freezing subgroups (vi). Data presented as individual data points and mean ± SEM. * significant differences CTR vs. PD, p<0.05; **p<0.01; #### main effect of sex p<0.0001.

Analyzing neuronal activation by PD and contextual fear memory retrieval revealed similar male-specific activation patterns for the medial part but not lateral part of the SuM. In sex-pooled groups, PD evoked neither a significant differences of cFos-positive cells in the medial SuM (Fig. 3Bi; n=14 CTR [one CTR male excluded due to insufficient representation of SuMM], n=18 PD, SuMM; Student’s T-test: T-test t(30)=1.685, p=0.1024) nor in the lateral SuM (Fig. 3Ci; n=14 CTR, n=17 PD [one male CTR and one male PD excluded due to insufficient representation of SuML], SuML; Student’s T-test: T-test t(29)=0.9953, p=0.3278). However, the two-way ANOVA revealed main sex effects for c-Fos in SuMM (Fig. 3Bii+iii; Male: n=8 CTR, n=7-8 PD; Female: n=6 CTR, n=10 PD; two-way ANOVA: sex: F(1,28)=9.395, p=0.0048; PD: F(1,28)=2.205, p=0.1488; interaction: F(1,28)=2.406 p=0.1321) and SuML (Fig. 3Cii+iii; two-way ANOVA: sex: F(1,27)=15.58, p=0.0005; PD: SuML F(1,27)=0.3400, p=0.5646; interaction: F(1,27)=1.008 p=0.3242). The separate analysis per sex demonstrated an increased c-Fos activation of the SUMM in males (Fig. 3Bii; n=8 CTR, n=8 PD; Student’s T-test: T(14)=2.823, p=0.014), while the SuML was not significantly activated (Fig. 3Cii; n=8 CTR, n=7 PD; Student’s T-test: T(13)=1.293, p=0.218). Again, female mice showed no change in SuM activation, neither in the medial (Fig. 3Biii; n=6 CTR, n=10 PD; Student’s T-test: T(14)=0.039, p=0.970) nor in the lateral portion (Fig. 3Ciii; n=6 CTR, n=10 PD; Student’s T-test: T(14)=0,269, p=0.792). The further analysis of female subgroups did not reveal an activation of SuM related to their freezing levels (Fig. 3Biv & 3Civ; one-way ANOVA: SuMM: F(2,7)=0.175, p=0.843; SUML: F(2,7)=0.123, p=0.886).

**Figure 3:**
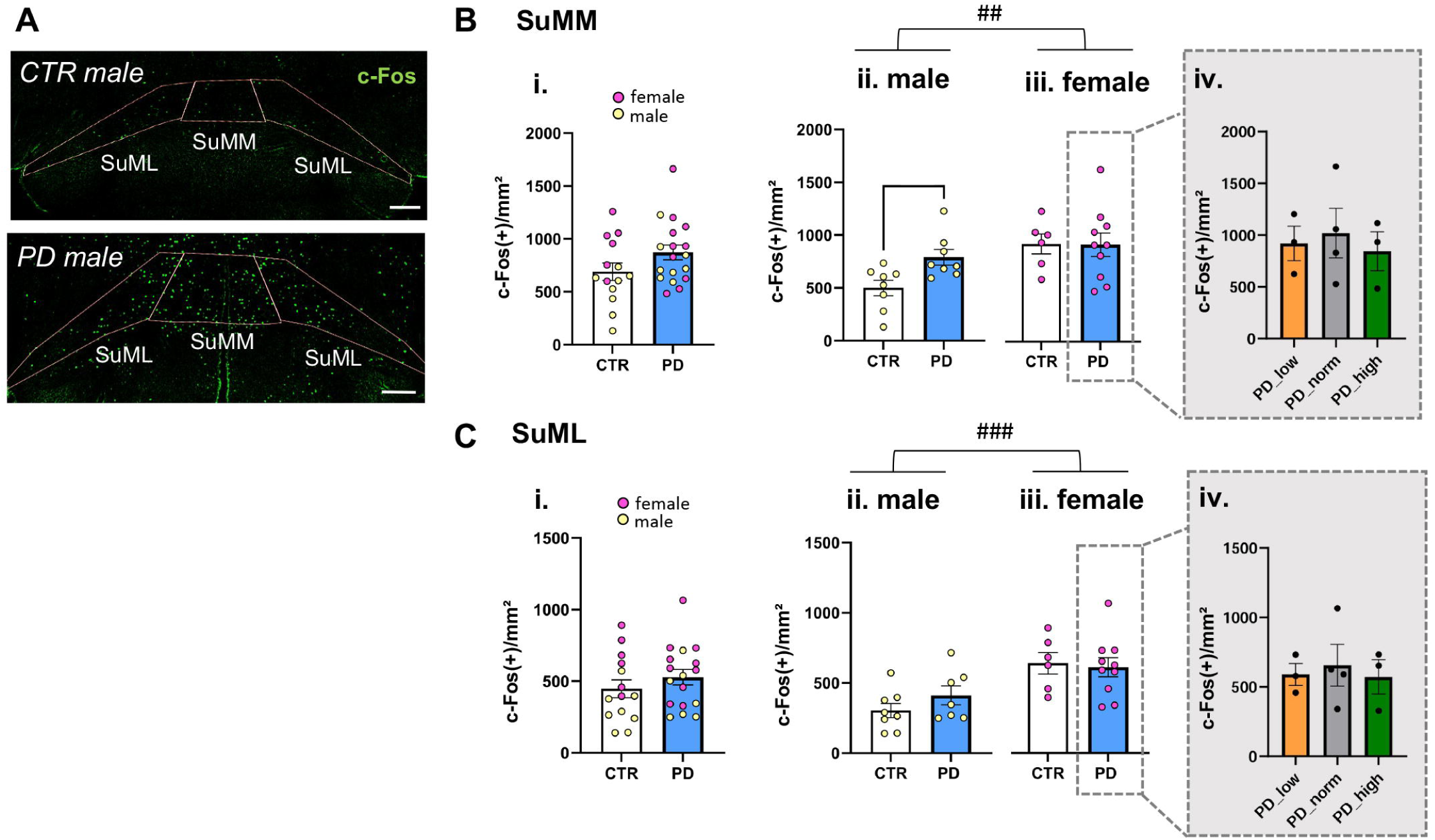
Phase delay (PD) overactivates the supramammilary nucleus (SuM) during fear memory retrieval in male mice. **(A)** Representative microphotographs of c-Fos immunolabelling (green) in the medial (SuMM) and lateral (SuML) SuM of male mice under control (CTR) and phase delay (PD) conditions, scale bar: 200 µm. **(B)** PD did not elicit alterations in SuMM activation when considering mice of both sexes (i), but two-way ANOVA revealed a main effect of sex. The separate analysis per sex showed an increased number of c-Fos(+) cells in male (iii), but not female mice after PD (iii). No differences were observed between female PD freezing subgroups (vi). **(C)** No change in c-Fos activation by PD of the SuML was observed for the whole cohort (i) or for sex-segregated groups (ii+iii), as well as when considering freezing subgroups in female mice (iv). Data presented as individual data points and mean ± SEM. * significant differences CTR vs. PD, p<0.05; ## main effect of sex p=0.01, ### p<0.001.

Within the dorsal hippocampus, a PD-effect was not evident on DG neuronal activation measured by c-Fos-positive cells on the hilus (Fig. 4Ai; n=15 CTR, n=18 PD; Mann-Whitney U-test: U=83, p=0.0620) or in the granule cell (GC) layer (Fig. 4Av; n=15 CTR, n=18 PD; Student’s T-test: T(31)=0.8181, p=0.4195). However, sex and PD effects were confirmed in the DG hilus (Fig. 4Aii+iii; two-way ANOVA on log10-transformed data: sex: F(1,29)=24.33, p<0.0001; PD: F(1,29)=4.635, p=0.0398; interaction: F(1,29)=1.200, p=0.2823) and in the DG GC a main sex effect was observed (Fig. 4Avi+vii; two-way ANOVA: sex: F(1,29)=16.85, p=0.0003; PD: F(1,29)=0.0857, p=0.7718; interaction: F(1,29)=1.714, p=0.2007). The separate analysis per sex demonstrated an increase of c-Fos positive cells by PD in the DG hilus of male mice (Fig. 4Aii; n=9 CTR, n=8 PD; Student’s T-test: T(15)=2.459, p=0.027), but not female mice (Fig. 4Aiii; n=6 CTR, n=10 PD; Student’s T-test: T(14)= 0.0583, p=0.954) and neither with respect to female freezing subgroups (Fig. 4Aiv; one-way ANOVA: F(2,7)=0.1698, p=0.847). No PD changes were observed within the GC of male (Fig. 4Avi; n=9 CTR, n=8 PD; Student’s T-test: T(15)=1.580, p=0.135) and female mice (Fig. 4Avii; n=6 CTR, n=10 PD; Student’s T-test: T(14)=0.573, p=0.576). However, the further analysis of female freezing subgroups showed a GC activation response (Fig. 4Aviii; one-way ANOVA: F(2,7)=7.744, p=0.017) with a significantly increased activation in the low-freezing females compared to the PD_norm group (comparable freezing levels to control group), as revealed by Tukey’s post hoc.

**Figure 4:**
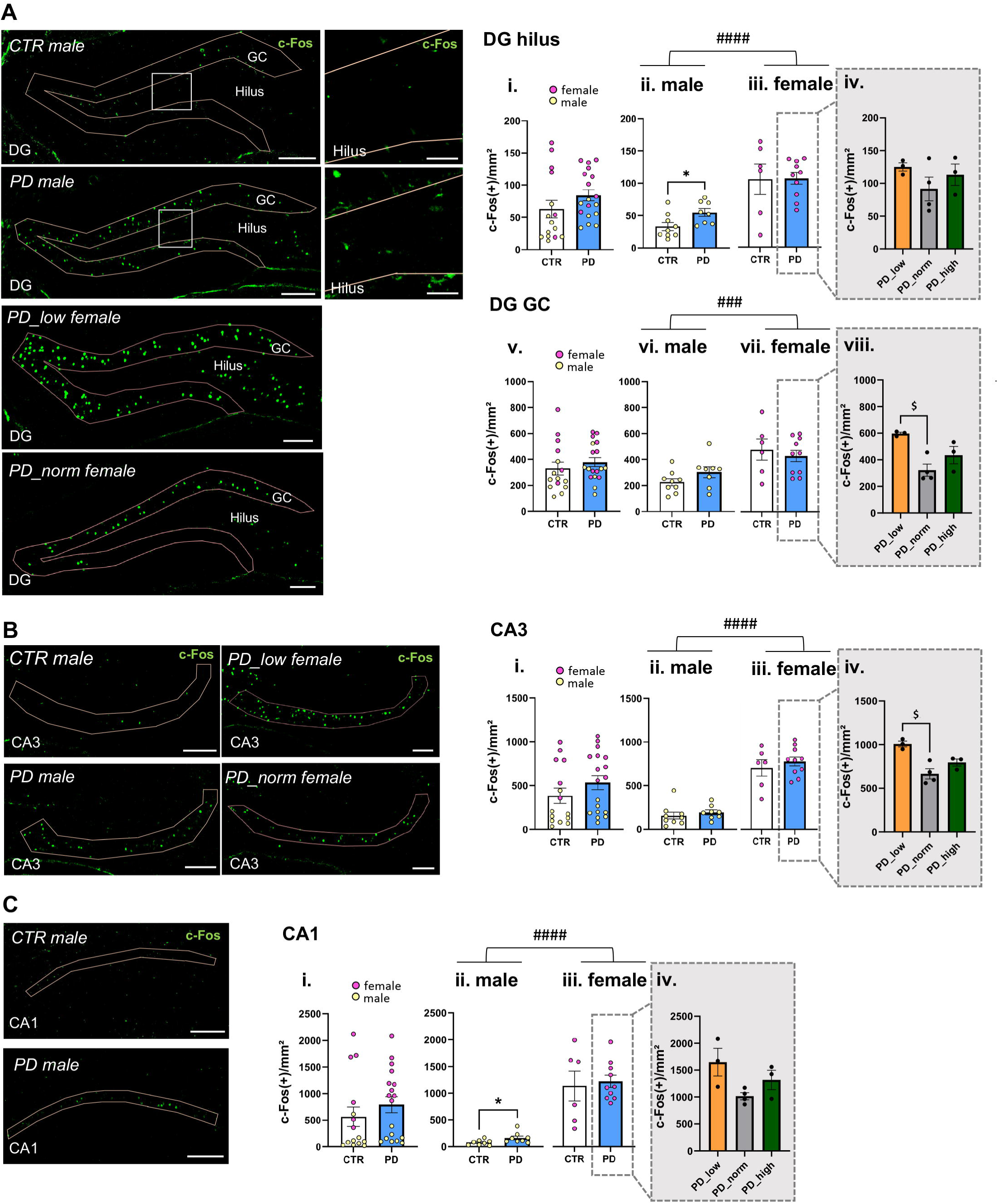
Phase delay (PD) overactivates the dorsal dentate gyrus (DG) during fear memory retrieval in male mice. **(A)** Representative microphotographs of c-Fos immunolabelling (green) in the dorsal dentate gyrus (DG) of control (CTR) and phase delay (PD) mice. Left scale bar: 200 µm; right: insert showing the hilus, scale bar: 50 µm. (i) Hilar cell activation was not altered by PD when analyzed across sexes, but two-way ANOVA demonstrates a main effect of sex and light. Separate analysis per sex indicates increased c-Fos–positive cell counts in the DG hilus of male (ii), but not female mice after PD (iii). No differences were observed between female PD freezing subgroups (vi). In the granule cell layer (GC) of the DG, PD did not shift cell activation when considering sex-pooled groups (v), but two-way ANOVA demonstrates a main effect of sex. The separate analysis per sex revealed no differences by PD in male (vi) or female mice (vii). However, within the females, the low freezing subgroup (PD_low) showed increased activation in the GC layer relative to mice with freezing levels comparable to the control group (PD_norm) (viii). **(B)** Representative microphotographs of c-Fos immunolabelling (green) in the dorsal CA3 region of CTR and PD mice, scale bar: 200 µm, and quantification of c-Fos(+) cells. No PD effect was detected for the cohort when pooling male and female mice (i), but two-way ANOVA revealed a main effect of sex. The separate analysis per sex revealed no differences by PD in male (ii) or female mice (iii). However, within the females, the low freezing subgroup (PD_low) showed increased activation in the GC layer compared to the PD_norm subgroup (iv). **(C)** Representative microphotographs of c-Fos immunolabelling (green) in the dorsal CA1 region of CTR and PD mice, scale bar: 200 µm. (i) Quantification of c-Fos(+) cells in the CA1 pyramidal layer did not show differences after PD when both sexes were analyzed combined (i), but the two-way ANOVA revealed a main effect of sex. The separate analysis per sex showed an increased activation by PD in male (ii), but not female mice (iii). No differences were observed between female PD freezing subgroups (vi). Data presented as individual data points and mean ± SEM. * significant differences CTR vs. PD, p<0.05; ### main effect of sex, p<0.001, #### p<0.0001; $ significant differences between female PD groups, p<0.05.

Neuronal activation was not shifted in the dorsal CA3 pyramidal layer following PD when comparing groups regardless of sex (Fig. 4Bi; n=15 CTR, n=18 PD, Mann-Whitney U-test: U=94, p=0.14448), but the two-way ANOVA revealed a main effect of sex (Fig. 4Bii+iii; males: n=9 CTR, n=8 PD; females: n=6 CTR, n=10 PD; two-way ANOVA on log10-transformed data: sex: F(1,29)=88.04, p<0.0001; PD: F(1,29)=2.193, p=0.1494; interaction: F(1,29)=0.4638, p=0.5013). The sex-specific analysis of PD effects did not reveal any differences in male (Fig. 4Bii; males: n=9 CTR, n=8 PD, Mann-Whitney U-test: U=23, p=0.236) or female mice (Fig. 4Biii: n=6 CTR, n=10 PD, Student’s T-test: T(14)= 0.764, p=0.458). The further analysis in female freezing response subgroups showed that CA3 neuronal activation differed significantly (Fig. 4Aiv; one-way ANOVA: F(2,7)=6.838, p=0.023), with increased neuronal activation in the low-freezing subgroup compared to the PD_norm (Tukey’s post hoc test, p=0.019).

In the dorsal CA1 pyramidal layer, PD did not shift neuronal activation when considering both sexes (Fig. 4Ci; n=14 CTR [one male Outlier removed], n=18 PD; Mann-Whitney U-test: U=95, p=0.1552), but again a main effect of sex was found (Fig. 4Cii+iii; males: n=8 CTR, n=8 PD; females: n=6 CTR, n=10 PD; two-way ANOVA on log10-transformed data: Sex F(1,29)=90.60, p<0.0001; PD: F(1,29)=2.784, p=0.1060; interaction: F(1,29)=0.5215, p=0.5215). Sex-specific comparisons showed that a higher activation of the CA1 pyramidal layer was observed in males (Fig. 4Cii; males: n=8 CTR, n=8 PD, Student’s T-test: T(14)=2.181, p=0.047), but not females (Fig. 4Ciii; n=6 CTR, n=10 PD, Student’s T-test: T(14)=0.333, p=0.744). Further analysis in female freezing subgroups did not reveal a differential neuronal activation in the CA1 layer (Fig. 4Civ; one-way ANOVA: F(2,7)=2.821, p=0.126).

Together, the circuit activation analysis suggests an activation of the SuM-DG circuit in male mice after PD exposure, together with an increased activation of orexinergic neurons in the LH. Nevertheless, DG-CA3 activity in females was related to their freezing response after PD.

### Chemogenetic overactivation of the dorsal dentate gyrus impairs fear memory expression and activates the supramammillary nucleus

To mimic the over-activation of the hilus of the DG after acute PD light shift, males received a chemogenetic activation of the dDG (DG+) during retrieval. DG overactivation led to a significant reduction of the time mice spent freezing (Fig. 5B; n=10 CTR, n=10 DG+; Student’s T-test: T(18)=3.792, p=0.001), thereby indeed mimicking the PD-induced phenotype. Investigating the activation of the DG, RNAScope for c-Fos mRNA confirmed the induction of neuronal activation in the DG hilus but also in DG GCs following our chemogenetic intervention (Fig. 5C+D; n=8 CTR, n=8-9 DG+; Mann-Whitney U-test: GC, U=1, p<0.001; hilus, U=1, p<0.001). Moreover, the activation of the dDG led to an overactivation of the CA3 and CA1 subregions of the hippocampus (Fig. S2; n=8 CTR, n=9 DG+, Mann-Whitney U-test: CA3, U=12, p=0.012; CA1, U=3, p<0.001).

**Figure 5:**
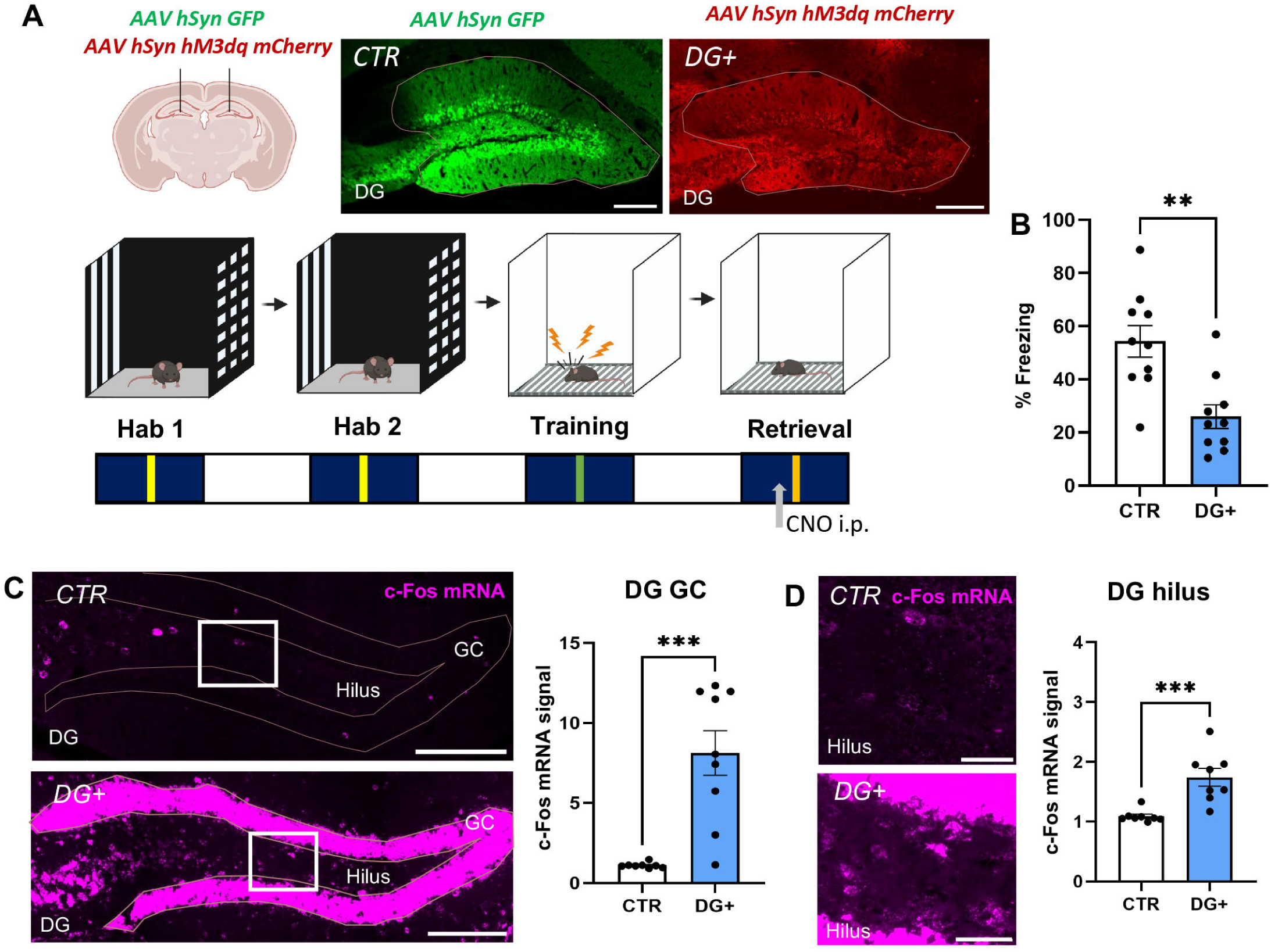
Chemogenetic overactivation of the dorsal dentate gyrus (DG) reduces freezing during contextual fear memory retrieval. **(A)** Male mice received injections of excitatory DREADD (DG+) or control vector (CTR) in the dorsal DG, representative microphotographs of viral expression patterns, scale bar: 250 µm. Two weeks later, contextual fear conditioning under standard 12h light/dark conditions took place during the dark active phase, with injection of CNO (1 mg/kg BW i.p., grey arrow) to activate the dorsal DG during retrieval of contextual fear memory. **(B)** DG chemogenetic activation decreased freezing compared to CTR. **(C)** Verification of DG overactivation by c-Fos immunolabelling, representative images of the DG total (left, scale bar 200 µm) and the hilar region (right, scale bar 50 µm) of CTR vs. DG+ mice. **(D)** Quantification of c-Fos intensities for the DG granule cell (GC) layer and cells in the hilus demonstrates overactivation of both subareas. Data presented as individual data points and mean ± SEM. ** significant differences CTR vs. DG+, p<0.01; ***p<0.001.

While the activation of orexinergic neurons in the LH was not altered after DG+ during retrieval (Fig. 6B; n=8 CTR, n=8 DG+; Student’s T-test: T(14)=1.512, p=0.153), an increased activation of the SuM was observed. Here, both the medial and the lateral subregions of the SuM, showed a higher expression of c-Fos mRNA after overactivation of the DG (Fig. 6D; n=8 CTR, n=8 DG+; SuMM, Student’s T-test: T(14)=2.295, p=0.038; SuML, Welch’s T-test: T(10.60)=2.701, p=0.021). The RNAScope approach further allowed the assessment of OxR1 mRNA expression in the SuM and the neuronal activation of OxR1-positive cells by their co-labeling with c-Fos mRNA. While the number of OxR1-postive cells was not altered by DG+ in the medial and lateral SuM subareas (Fig. 6E; n=8 CTR, n=8 DG+; SuMM, Student’s T-test: T(14)=0.818, p=0.427; SuML, Student’s T-test: T(14)=0.076, p=0.941), the portion of OxR1-postive cells activated by c-Fos did increase by DG+ (Fig. 6F; n=8 CTR, n=8 DG+; SuMM, Student’s T-test: T(14)=2.347, p=0.034; SuML, Student’s T-test: T(14)=3.007, p=0.009).

**Figure 6:**
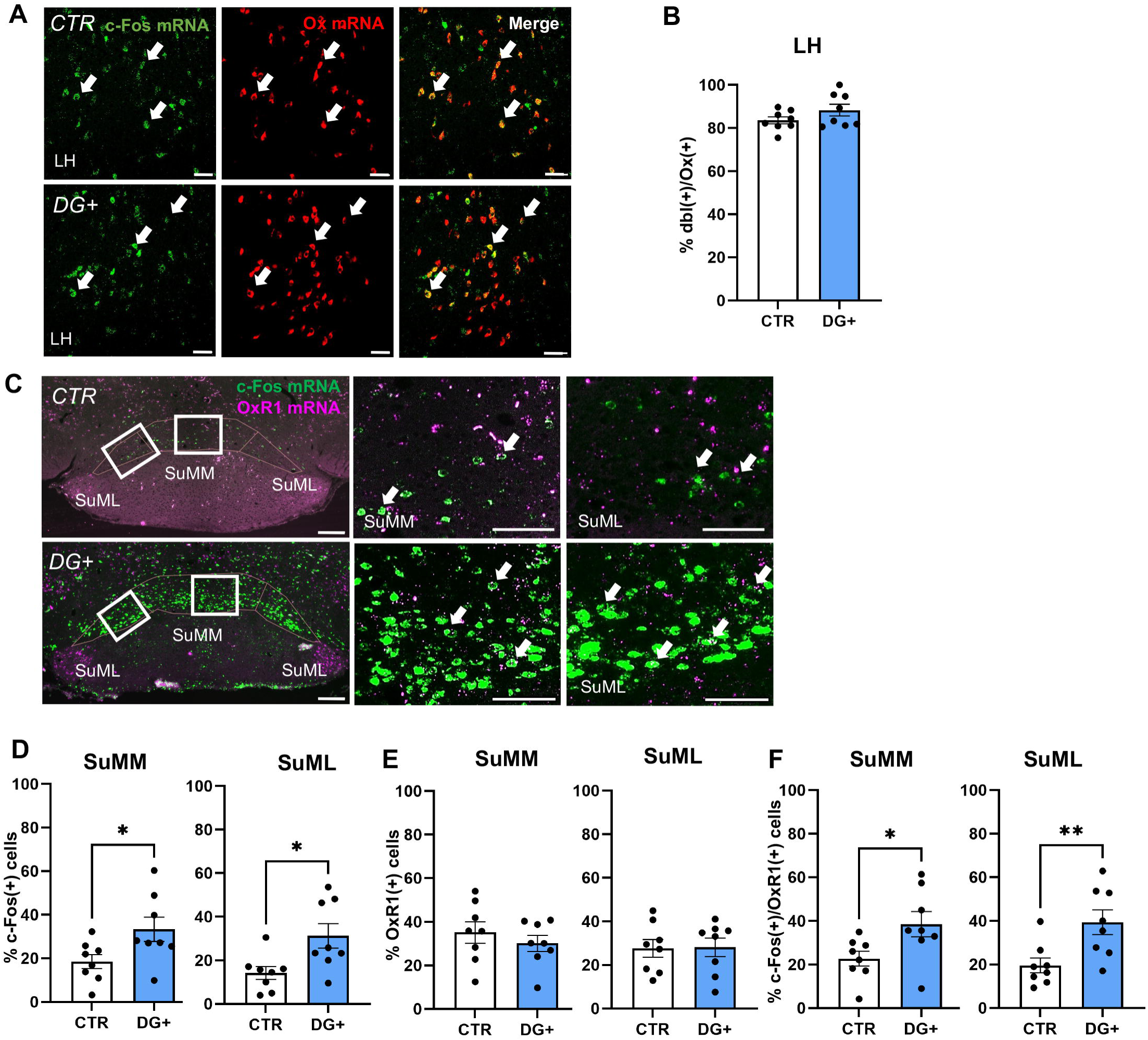
Chemogenetic overactivation of the dorsal dentate gyrus (DG) during contextual fear memory retrieval activates the supramammillary nucleus (SuM). **(A)** Representative microphotographs of c-Fos (green), orexin (Ox, red), and double (merged, yellow) immunolabelling in the lateral hypothalamus (LH; white arrow), scale bar: 50 µm. (B) DREADD activation of the DG during retrieval did not affect the proportion of c-Fos(+) cells within the orexinergic population in the LH (dbl, c-Fos(+)-Ox(+)). (C) Representative microphotographs of c-Fos in situ hybridisation (green) in the medial (SuMM) and lateral (SuML) SuM, scale bar: 200 µm. **(D)** Chemogenetic DREADD activation of the DG during retrieval significantly increased the proportion of c-Fos(+) cells in the SuMM and SuML. **(E)** No differences in OxR1 expression in the SuM subregions. **(F)** Chemogenetic activation of the DG increased the percentage of c-Fos(+) cells within the OxR1(+) population. Data presented as individual data points and mean ± SEM. * significant differences CTR vs. DG+, p<0.05; **p<0.01.

Together, the overactivation of the DG mimics the PD-induced phenotype and recapitulates the activation of the DG-SuM circuit, but not of orexinergic neurons in the LH.

### Chemogenetic overactivation of the supramammillary nucleus impairs fear memory expression and activates the orexinergic neurons in the lateral hypothalamus

Next, the activation of the SuM during fear memory retrieval was tested as well to mimic its PD-induced increase in c-Fos. The chemogenetically activated SuM led again to a significant reduction of freezing levels during retrieval compared with controls (Fig. 7B; n=6 CTR, n=6 SuM+; Mann-Whitney U-test: U=2, p=0.009). The efficacy of local SuM DREADD activation was confirmed by an increased density of c-Fos immunopositive cells in the medial and lateral SuM subareas (Fig. 7D; n=6 CTR, n=6 SuM+; SuMM, Welch’s T-test: T(5.212)=8.039, p<0.001; SuML, Mann-Whitney U-test: U=0, p=0.002). In contrast to DG+, orexinergic neurons in the LH were activated as well by SuM+, as assessed by an increase in c-Fos-positive of OxA-positive cells (Fig. 8B; n=6 CTR, n=5 SuM+; Welch’s T-test: T(7.118)=2.690, p=0.031).

**Figure 7:**
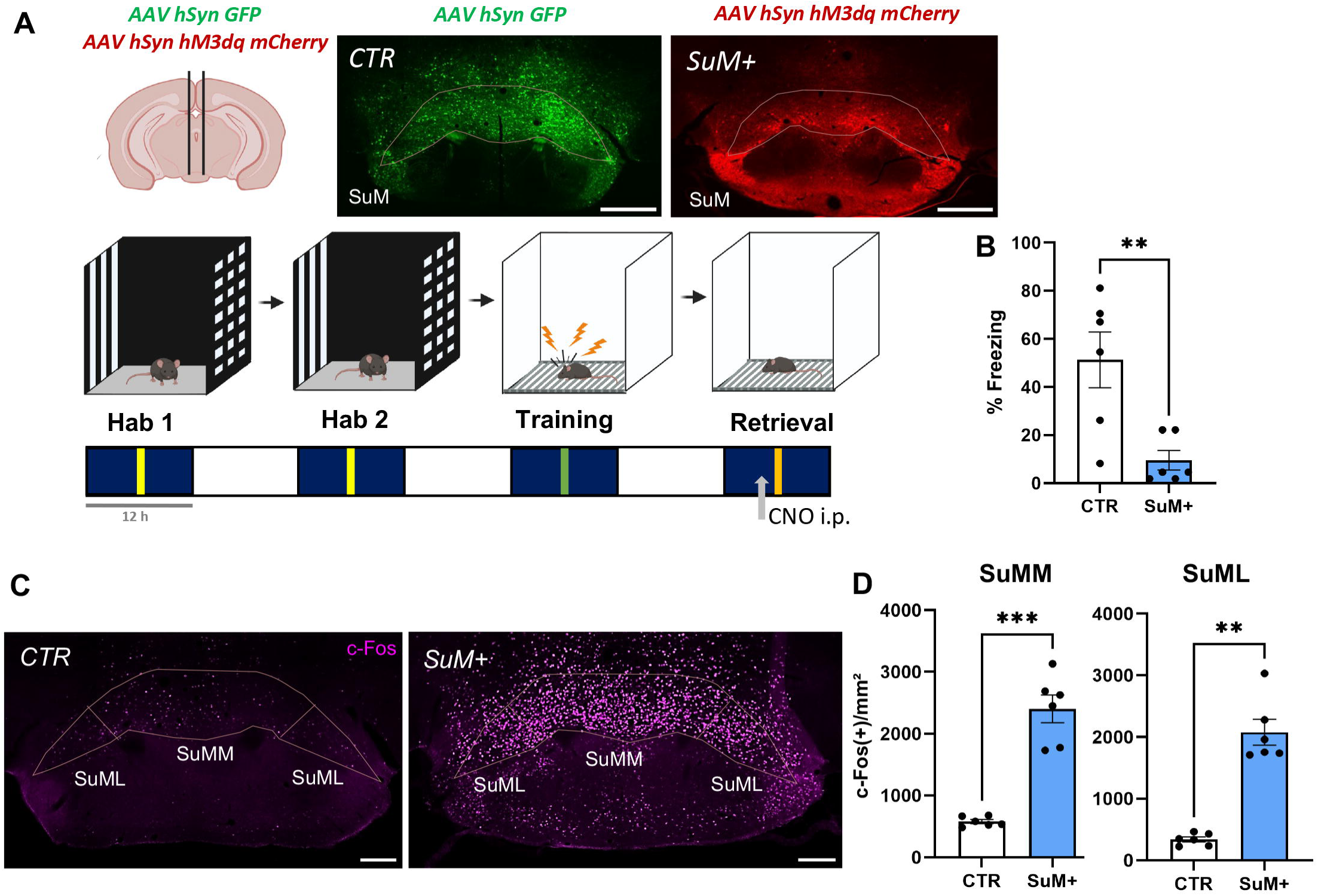
Chemogenetic overactivation of the supramammillary nucleus (SuM) reduces freezing during contextual fear memory retrieval. **(A)** Male mice received injections of excitatory DREADD (SuM+) or control vector (CTR) in the SuM, representative microphotographs of viral expression patterns, scale bar: 500 µm. Two weeks later, contextual fear conditioning under standard 12 h light/dark conditions took place during the dark active phase, with injection of CNO (5 mg/kg BW i.p., grey arrow) to activate the SuM during retrieval of contextual fear memory. **(B)** SuM chemogenetic activation decreased freezing compared to the CTR. **(C)** Representative microphotographs for c-Fos immunolabelling, scale bar: 200 µm. **(D)** Quantification of c-Fos(+) cells in the SuM verifies the chemogenetic overactivation of its medial and lateral subregions. Data presented as individual data points and mean ± SEM. ** significant differences CTR vs. SuM+, p<0.01; ***p<0.001.

**Figure 8:**
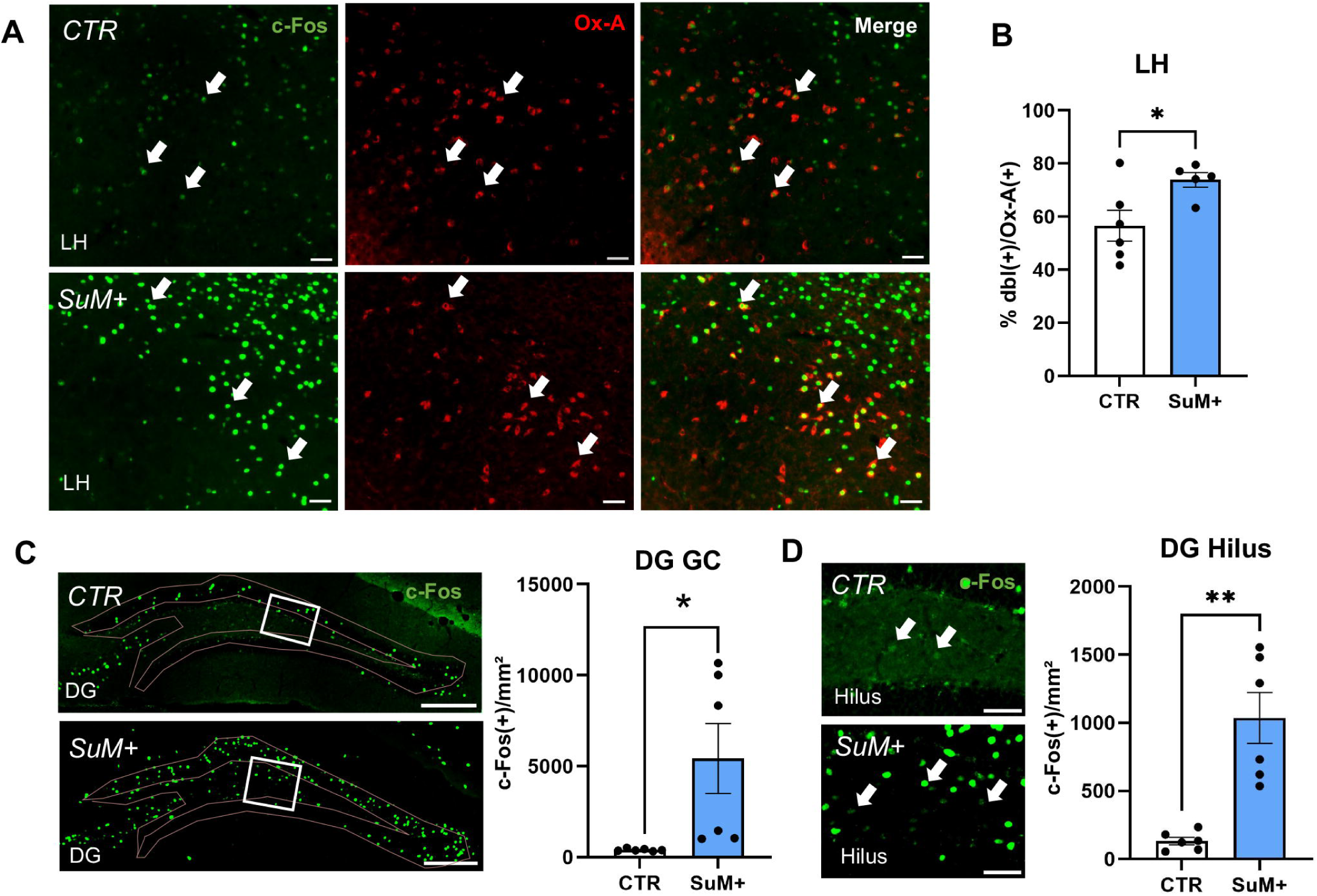
Chemogenetic overactivation of the supramammillary nucleus (SuM) activates orexinergic neurons in the lateral hypothalamus (LH) and hilar cells in the dorsal dentate gyrus (DG) during contextual fear memory retrieval. **(A)** Representative microphotographs of c-Fos (green) and orexin-A (Ox-A, red) immunolabelling after overactivation of the SuM, scale bar: 50 µm. (B) Quantification of double-labelled neurons revealed an induction of orexinergic neurons. (C) Representative microphotographs of c-Fos (green) immunolabelling in the dorsal DG after overactivation of the SuM, scale bar: 200 µm, and quantification in the DG revealing activation of the DG granule cells (GC). (D) Representative microphotographs of c-Fos (green) immunolabelling in the hilus of the dorsal DG after overactivation of the SuM, scale bar: 50 µm. SuM overactivation increased the proportion of hilar cells activated by c-Fos. Data presented as individual data points and mean ± SEM. * significant differences CTR vs. SuM+, p<0.05; **p<0.01.

Moreover, chemogenetic SuM activation during retrieval increased the neuronal activation in the DG GC and hilus (Fig. 8C+D; n=6 CTR, n=6 SuM+; GC, Mann-Whitney U-test: U=0, p=0.0022; Hilus, Welch’s T-test: T(5,217)=4.784, p=0.0044). Such an increased activation was observed along the trisynaptic hippocampal pathway as well, in the CA3 and CA1 pyramidal cell layer (Fig. S3; n=6 CTR, n=6 SuM+; CA3, Welch’s T-test: T(5,254)=3.403, p=0.018; CA1, Student’s T-test: T(10)=6.955, p<0.0001). These data demonstrate that the chemogenetic activation of the SuM during retrieval leads to reduced freezing, in association with an overactivation of the SuM-DG pathway and of orexinergic neurons in the LH.

## Discussion

In this study, we demonstrated that an acute delay of the dark phase after CFC impaired fear expression in male, but not in female mice. This behavioral deficit was accompanied by an increased expression of the neuronal activity marker c-Fos in the hilus of the DG and in the medial subregion of the SuM, suggesting an overactivation of the SuM-DG circuit, as well as in orexin-positive neurons in the LH. No significant shifts in neuronal activity were detectable in female mice, mirroring their behavioral phenotype. Chemogenetic overactivation of DG or SuM during retrieval mimicked the PD-induced phenotype and confirmed SuM-DG interactions. Moreover, chemogenetic SuM activation recruited more orexinergic cells. Collectively, these findings suggest that PD following aversive memory formation activates orexinergic neurons and the SuM-DG circuit in a male mice, leading to reduced fear expression.

Acute shifts of the LD phase have been shown to alter fear memory acquisition and expression in male mice: 12 h PD of the dark phase before CFC training impaired freezing, as well as a 6 h PD or PA directly after training (Loh et al., 2010). While Loh and colleagues report a stronger impact of PA on fear expression, we observed a significant effect of PD, similar to observations on cued fear memory reconsolidation (Clark et al., 2020). These discrepancies may relate to differences in training timing. While Loh et al. (2010) fear-conditioned mice during the light phase, we trained and tested the animals during the active, dark phase. Additionally, PA by Loh et al. promoted an earlier onset of the dark phase, while PD in our hands prolonged this phase. Thus, a net prolongation of the dark phase after CFC training may induce deficits in fear expression overall. Although the total amount of sleep was not changed, they reported changes in the distribution of resting episodes after the phase shifts. Acute phase shift during our study may therefore alter sleep patterns after CFC training. In humans and rodents, REM sleep is required for memory consolidation (Pedrazzoli et al., 2004; Xia & Storm, 2017), with the early consolidation phase (4-6 h post-training) being particularly sensitive to enforced sleep deprivation (SD) (Graves et al., 2003; Ishikawa et al., 2014). In our design, the prolonged dark phase aligns with a late consolidation window (8-10 h), typically not sleep-dependent (Graves et al., 2003; Ishikawa et al., 2014). However, fragmented sleep may influence fear memory expression, as it has been demonstrated in a mouse model of reduced cholinergic transmission (Queiroz et al., 2013) and after shortening the LD phases for at least two weeks (Lee et al., 2016).

Male rodents appear more susceptible to the effect of altered sleep patterns on fear memory. SD after CFC disrupted fear extinction in male, but not female rats (Hunter, 2018) and chronic SD impairs fear memory more severely in male than female mice (Sajadi et al., 2025). Brief episodes of bright light exposure during the dark phase induced sex-specific effects as well, with reduced recognition memory and increased locomotion in male mice, but distinct estrous cycle-dependent effects in females (Datta et al., 2019). In our study, acute PD induced no differences in fear expression in females. Freezing levels were generally higher in females, consistent with previous studies (Botterill et al., 2021), which may affect the detection of group differences. Moreover, since our group sizes would allow to detect only large effects, adaptations to the protocol and using larger cohorts of females may allow to a more in-depth analysis of behavioral and circuit adaptations by PD in females. In this line, even with our small cohort, an individual profiling revealed a heterogeneous freezing response in females. In females with a low freezing response after PD, an increased activation of the dDG GC and the CA3 region was observed, underlining the relevance of these regions in mediating fear expression. No other PD-induced circuit changes were observed, suggesting a compensatory mechanism in females that promotes network stability against acute phase shift. Further studies are needed to investigate this effect and to identify sex-dependent mediators for network stability in females. While the estrous phase was not assessed to avoid additional vaginal smear sampling stress, sex- and estrous cycle-dependent expression of OxRs and pre-pro-orexin (Gao & Horvath, 2021; Silveyra et al., 2007) are compelling candidates, since sexually dimorphic expression of OxR1 and OxR2 in the amygdala modulates sex-specific responses to social stress (Yaeger et al., 2026).

In males, increased c-Fos in orexin-positive LH neurons was observed after PD and fear memory recall, similar to observations after SD (Estabrooke et al., 2001; Modirrousta et al., 2005). A recent study further demonstrated that the pharmacological manipulation of OxRs during a reward memory paradigm was effective only in sleep-deprived animals (Almeida Rojo et al., 2025), suggesting that the orexinergic system mediates internal states to support memory performance under sleep loss. Receptor antagonist studies support the modulatory role of the orexinergic system on fear memory: systemic blockage of OxR1 reduced the expression of contextual fear in rats (Wang et al., 2017), with local interventions suggesting OxR1 action on locus coeruleus-lateral amygdala projections (Soya et al., 2017) and within the prelimbic cortex (Oliveira et al., 2025). OxR1 is expressed in the DG as well, with a higher density in the hilus than OxR2 (Krause et al., 2024), and has been linked to conditioned place preference (Guo et al., 2016), consolidation of passive avoidance and spatial memory (Akbari et al., 2007, 2008), supporting a role for orexin inputs to the DG in memory.

The role of the DG in CFC is well established. Optogenetic inhibition demonstrated a contribution of the DG to consolidation, retrieval, and extinction of contextual fear memories (Bernier et al., 2017). In epilepsy models with states of DG overactivation, contextual fear memory is impaired as well (Lima et al., 2016). Our DG chemogenetic stimulation may limit the precise reactivation of contextual engrams formed in the DG required for successful contextual fear retrieval (Chua & Tan, 2017; Cui et al., 2024; Iwasaki & Ikegaya, 2021; Wilmot et al., 2019). Hilar interneurons additionally modulate DG GC activity, thereby shaping contextual fear engrams (Raza et al., 2017) and fear expression (Comeras et al., 2021). DG chemogenetic activation during retrieval increased c-Fos in the CA3 and CA1 and in the SuM, indicating propagation along the trisynaptic hippocampal pathway and feedback to the SuM. This feedback may include indirect pathways, for example through the medial and lateral septum or medial prefrontal cortex (Pan and McNaughton, 2004; Parent et al., 2009), and a further modulation by long-range projections of CA1 somatostatin-positive interneurons (Kinney et al., 2025). Importantly, OxR1-positive neurons in the SuM were recruited during contextual fear memory in the control group, with further increase in c-Fos activation under DG overactivation, suggesting that fear-responsive SuM neurons are susceptible to orexinergic modulation.

The SuM is a well-established modulator of DG activity (Hashimotodani et al., 2018). SuM-to-DG projecting neurons co-release GABA and glutamate (Ajibola et al., 2021), with an excitatory net effect on GCs (Tabuchi et al., 2022) and may prime plasticity at DG synapses (Carre & Harley, 1991; Ito et al., 2018; Nakanishi et al., 2001). Projection-specific chemogenetic silencing of SuM-to-dDG neurons decreases freezing, while their optogenetic activation increases it in a CFC task (Luo et al., 2025). Our global SuM chemogenetic stimulation before fear memory retrieval reduced freezing, contradicting the projection-specific results. However, global SuM excitation may engage multiple downstream targets that contribute to fear expression, such as the LH, cortex, thalamus, brainstem, and medial septum (Pan & McNaughton, 2004). Additionally, excitatory projections from the SuM reach interneurons in the dorsal CA3 and CA1 subregions (Jiang et al., 2024; M. Li et al., 2023), but in our hands chemogenetic activation of the SuM increased c-Fos counts in the pyramidal cell layer of CA3 and CA1 and in DG GCs, suggesting that SuM stimulation enhanced the DG output propagating through the trisynaptic hippocampal circuit rather than triggering CA1 and CA3 interneurons.

Additionally, SuM overactivation induced a greater proportion of c-Fos-positive orexinergic cells. The LH and SuM are interconnected bidirectionally (Peyron et al., 1998; Plaisier et al., 2020). Since SuM neurons recruited during fear memory recall express OxR1, they are susceptible to orexinergic modulation. In addition, increased activation of SuM neurons as well as orexinergic neurons in the LH during PD could potentially converge onto the DG synapses (Hashimotodani et al., 2018; Peyron et al., 1998; Vertes, 2015), thereby modulating hilar-GC microcircuits.

## Conclusion & Future Directions

Our results identify a circuit activated by fear memory retrieval after acute PD that involves orexinergic neurons in the LH, the SuM and the DG in male mice (Fig. 9). Chemogenetic activation supports the DG and the SuM as main drivers for reducing fear expression. Pharmacological orexinergic interventions may help to dissect the modulation of the SuM-DG circuit further and may uncover compensatory mechanisms observed in female mice. Whether altered sleep patterns after PD may drive orexinergic activation warrants future studies. Due to the high prevalence of circadian misalignment and sleep disruption in our modern societies and the growing interest in using orexinergic-modulating drugs for the treatment of memory-related disorders such as Alzheimer’s (Kourosh-Arami et al., 2025; Lucey et al., 2023; Moline et al., 2021; Ragsdale et al., 2025), a detailed understanding of orexinergic effects on memory circuits has direct translational relevance and would help to improve treatment schemes.

**Figure 9:**
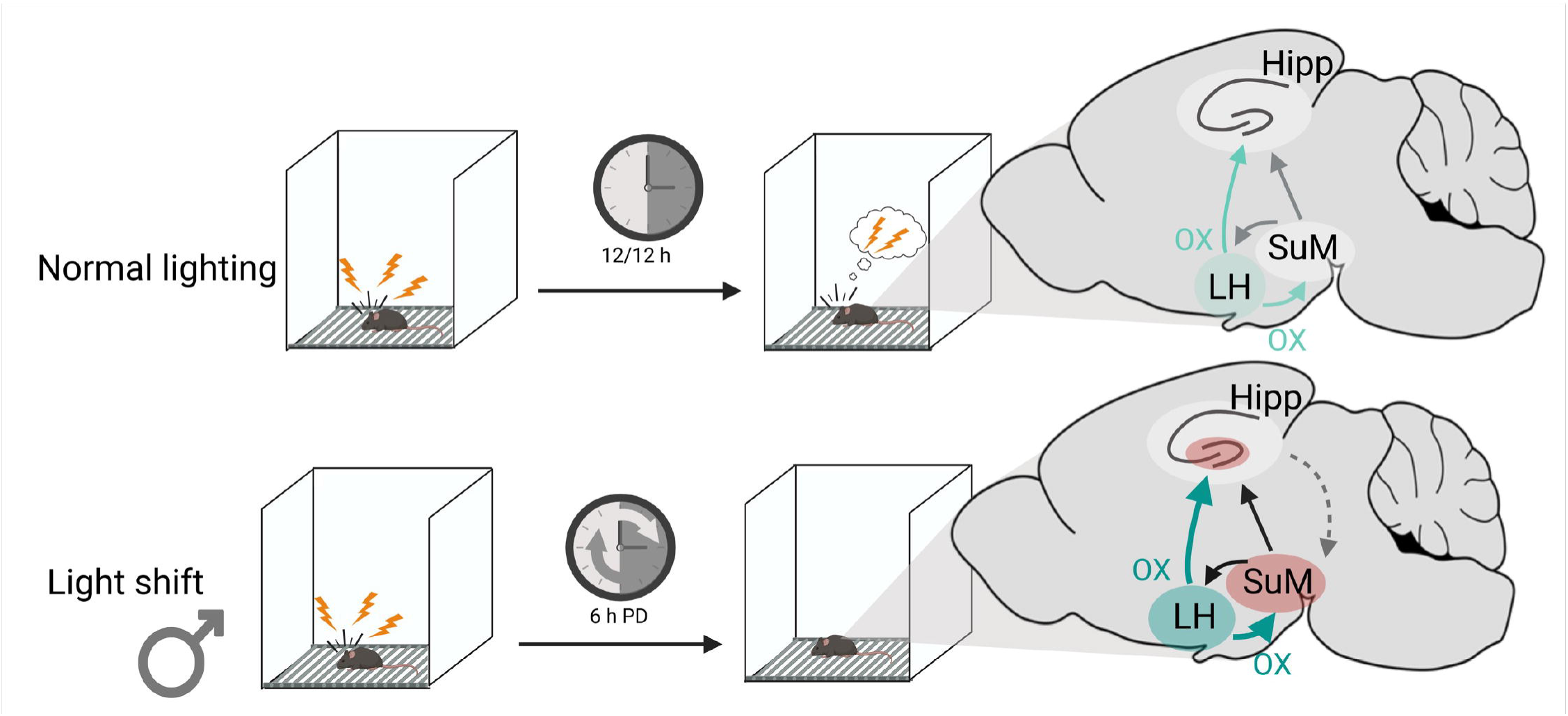
Summary graph for the proposed fear-circuit elements activated by acute phase delay (PD). **(A)** Recall of contextual fear memory during the dark phase activates the dorsal hippocampus, the supramammillary nucleus (SuM), and orexinergic neurons in the lateral hypothalamus (LH). **(B)** After a 6 h PD of the dark phase following fear conditioning training, male mice show reduced fear expression, which is associated with an overactivation of the SuM-DG circuit and increased activation of orexinergic neurons in the LH. In female mice, neither the behavioral response nor the circuit activation is observed after PD. Created in BioRender. Chirich, L. (2026) https://BioRender.com/be0vmjk

## Author Contributions

LCB: Conceptualization, Methodology, Formal analysis, Investigation, Writing – original draft, Writing – review & editing.

HG: Methodology, Formal analysis, Investigation, Writing – original draft, Writing – review & editing.

AA: Conceptualization, Formal analysis, Resources, Funding acquisition, Writing – original draft, Writing – review & editing.

## Acknowledgement

This work was supported by a grant from the German Research Foundation (Project-ID 425899996 – CRC 1436 to Anne Albrecht) and from the Center for Behavioural Brain Sciences Magdeburg - CBBS funded by the European funds for regional development (EFRE, Funding Nr ZS/2016/04/78113). We thank Annika Lenuweit, Romina Wolter and Leon Joris Schmidt for excellent technical assistance and the team of the central animal facility of the medical faculty of the Otto-von-Guericke University for excellent animal care.

