## Supplementary material for "Subcortical-hippocampal circuits for mediating impaired contextual fear memory after an acute shift of the light/dark phase": S1, S2, S3

### Supplementary information

**Supplementary Figure S1:** Acute phase advance (PA) does not change fear memory expression in male mice.

**Supplementary Figure S2:** Behavioural profiling of female mice after phase delay (PD) reveals hippocampal activation linked to freezing expression.

**Supplementary Figure S3:** c-Fos signal in the CA3 and the CA1 layer during retrieval under chemogenetic dentate gyrus (DG) activation

**Supplementary Figure S4:** c-Fos signal in the CA3 and the CA1 layer during retrieval under chemogenetic supramammillary nucleus (SuM) overactivation.

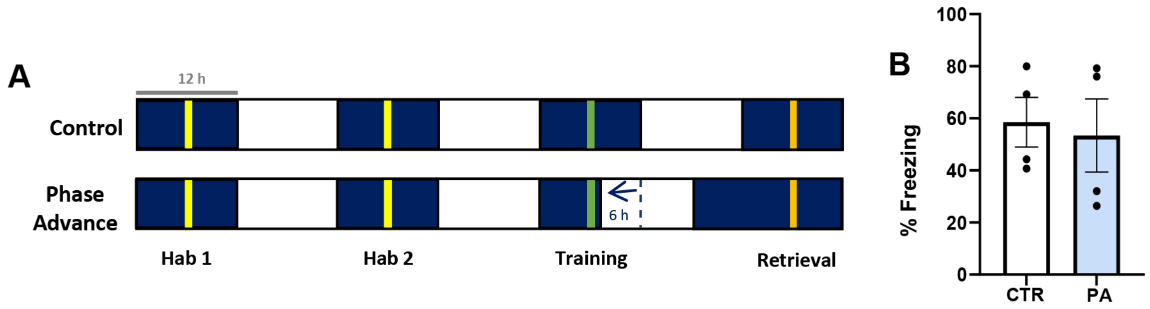


**Fig. S1: Acute phase advance (PA) does not change fear memory expression in male mice. (A)** Schematic timeline of contextual fear conditioning protocol with a 6-h phase advance (PA), resulting in earlier onset of the light phase after training, compared to control (CTR) light/ dark conditions. **(B)** Freezing was not significantly affected by PA. Data presented as individual data points and mean ± SEM.

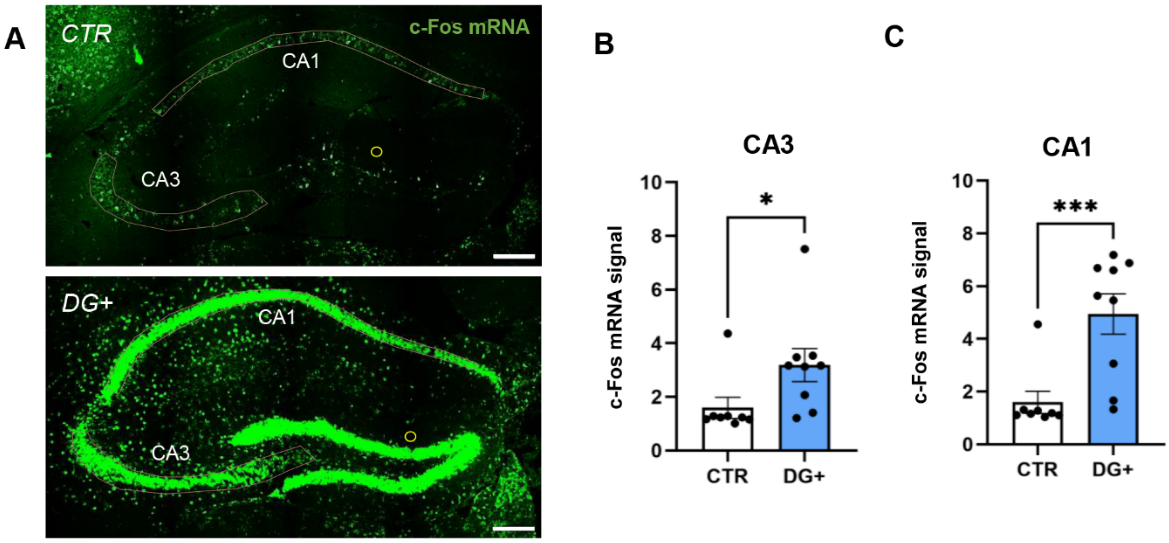


**Fig. S2: c-Fos signal in the CA3 and the CA1 layer during retrieval under chemogenetic dentate gyrus (DG) activation. (A)** Representative microphotographs of RNAScope fluorescence in situ hybridization for c-Fos mRNA (green) in the dorsal hippocampus, scale bar: 200 µm. **(B)** Hippocampal activation was calculated as the intensity of c-Fos mRNA normalized to the background signal (yellow circle). The normalized c-Fos mRNA signal was increased in the CA3 and **(C)** the CA1 region of the dorsal hippocampus following chemogenetic overactivation of the DG (DG+) during contextual fear memory retrieval compared to control animals (CTR). Data presented as individual data points and mean ± SEM. * significant differences CTR vs. DG+, $p<0.05$; ***$p<0.001$.

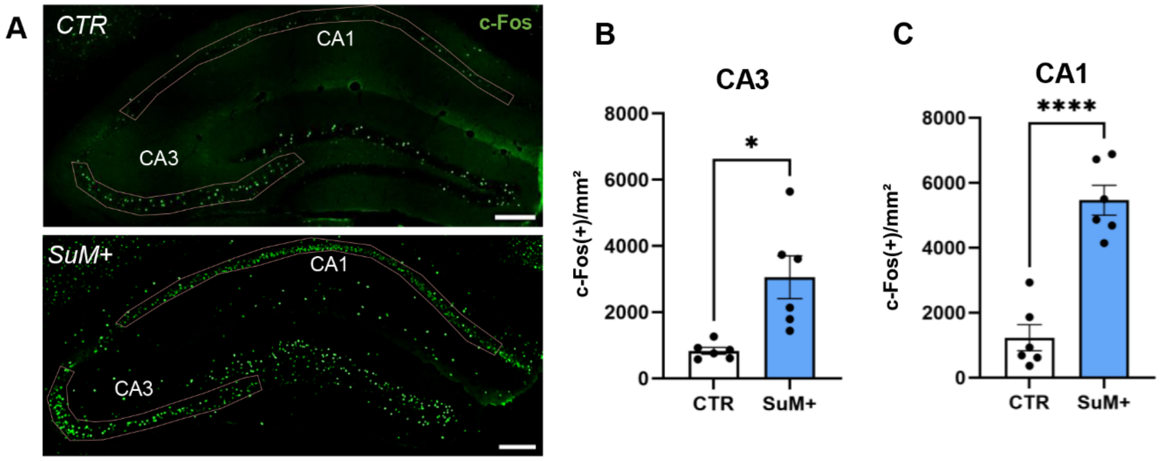


**Fig. S3: c-Fos signal in the CA3 and the CA1 layer during retrieval under chemogenetic supramammillary nucleus (SuM) activation. (A)** Representative microphotographs for c-Fos (green) immunolabelling in the dorsal hippocampus, scale bar: 200 µm. **(B)** Chemogenetic SuM overactivation (SuM+) during retrieval significantly increased the density of c-Fos(+) cells in the CA3 and **(C)** in the CA1 pyramidal layer of the dorsal hippocampus compared to control (CTR) animals. Data presented as individual data points and mean ± SEM. * significant differences CTR vs. SuM+, $p<0.05$; ****$p<0.0001$.
